# On the necessity of recurrent processing during object recognition: it depends on the need for scene segmentation

**DOI:** 10.1101/2020.11.11.377655

**Authors:** Noor Seijdel, Jessica Loke, Ron van de Klundert, Matthew van der Meer, Eva Quispel, Simon van Gaal, Edward H.F. de Haan, H. Steven Scholte

## Abstract

While feed-forward activity may suffice for recognizing objects in isolation, additional visual operations that aid object recognition might be needed for real-world scenes. One such additional operation is figure-ground segmentation; extracting the relevant features and locations of the target object while ignoring irrelevant features. In this study of 60 participants, we show objects on backgrounds of increasing complexity to investigate whether recurrent computations are increasingly important for segmenting objects from more complex backgrounds. Three lines of evidence show that recurrent processing is critical for recognition of objects embedded in complex scenes. First, behavioral results indicated a greater reduction in performance after masking objects presented on more complex backgrounds; with the degree of impairment increasing with increasing background complexity. Second, electroencephalography (EEG) measurements showed clear differences in the evoked response potentials (ERPs) between conditions around 200ms - a time point beyond feed-forward activity and object decoding based on the EEG signal indicated later decoding onsets for objects embedded in more complex backgrounds. Third, Deep Convolutional Neural Network performance confirmed this interpretation; feed-forward and less deep networks showed a higher degree of impairment in recognition for objects in complex backgrounds compared to recurrent and deeper networks. Together, these results support the notion that recurrent computations drive figure-ground segmentation of objects in complex scenes.

## Introduction

The efficiency and speed of the human visual system during object categorization suggests that a feed-forward sweep of visual information processing is sufficient for successful recognition (VanRullen and Thorpe, 2002). For example, when presented with objects in a rapid serial visual presentation task (RSVP; (Potter and Levy, 1969), or during rapid visual categorization (Thorpe et al., 1996), human subjects could still successfully recognize these objects, with EEG measurements showing robust object-selective activity within 150 ms after object presentation (VanRullen and Thorpe, 2001). Given that there is substantial evidence for the involvement of recurrent processing in figure–ground segmentation (Lamme and Roelfsema, 2000; Scholte et al., 2008; Wokke et al., 2012), this seems inconsistent with recognition processes that rely on explicit encoding of spatial relationships between parts and suggest instead that rapid recognition may rely on the detection of an ‘unbound’ collection of image features (Crouzet and Serre, 2011).

Recently, a multitude of studies have reconciled these seemingly inconsistent findings by indicating that recurrent processes might be employed adaptively, depending on the visual input: while feed-forward activity might suffice for simple scenes with isolated objects, more complex scenes or more challenging conditions (e.g. objects that are occluded or degraded), may need additional visual operations (‘routines’) requiring recurrent computations (Groen et al., 2018; Tang et al., 2018; Kar et al., 2019; Rajaei et al., 2019; Seijdel et al., 2020). For objects in isolation, or very simple scenes, rapid recognition may thus rely on a coarse and unsegmented feed-forward representation (Crouzet and Serre, 2011), while for more cluttered images recognition may require explicit encoding of spatial relationships between parts. In other words, for those images, extra visual operations to group parts of the object, and to segment this object (‘figure’) from its background might be needed.

Several studies have already shown that the ‘segmentability’ of a natural scene might influence the degree of recurrent processing. For example, Koivisto, Kastrati & Revuonso reported that masking, a technique shown to affect mainly recurrent but not feed-forward processing (Fahrenfort et al., 2007), was more effective for objects that were rated as being ‘difficult to segregate’ (Koivisto et al., 2014). Also in a more recent study we showed that natural scene complexity, providing information about the ‘segmentability’ of a scene, modulates the degree of feedback activity in the brain (Groen et al., 2018). However, both studies did not test for effects of segmentation explicitly and used natural scenes that were uncontrolled and in which complexity could correlate with other contextual factors in the scene. Therefore, we here systematically investigated whether scene complexity influenced the extent of recurrent processing during object recognition. To this end, participants performed an object recognition task with objects embedded in backgrounds of different complexity (Figure 1), indexed by two biologically plausible measures: the spatial coherence (SC) and contrast energy (CE) (Ghebreab et al., 2009; Scholte et al., 2009; Groen et al., 2013). Using these ‘hybrid’ stimuli, we combine relevant features of objects in natural scenes, embedded in well controlled backgrounds of different complexity.

**Figure 1.**
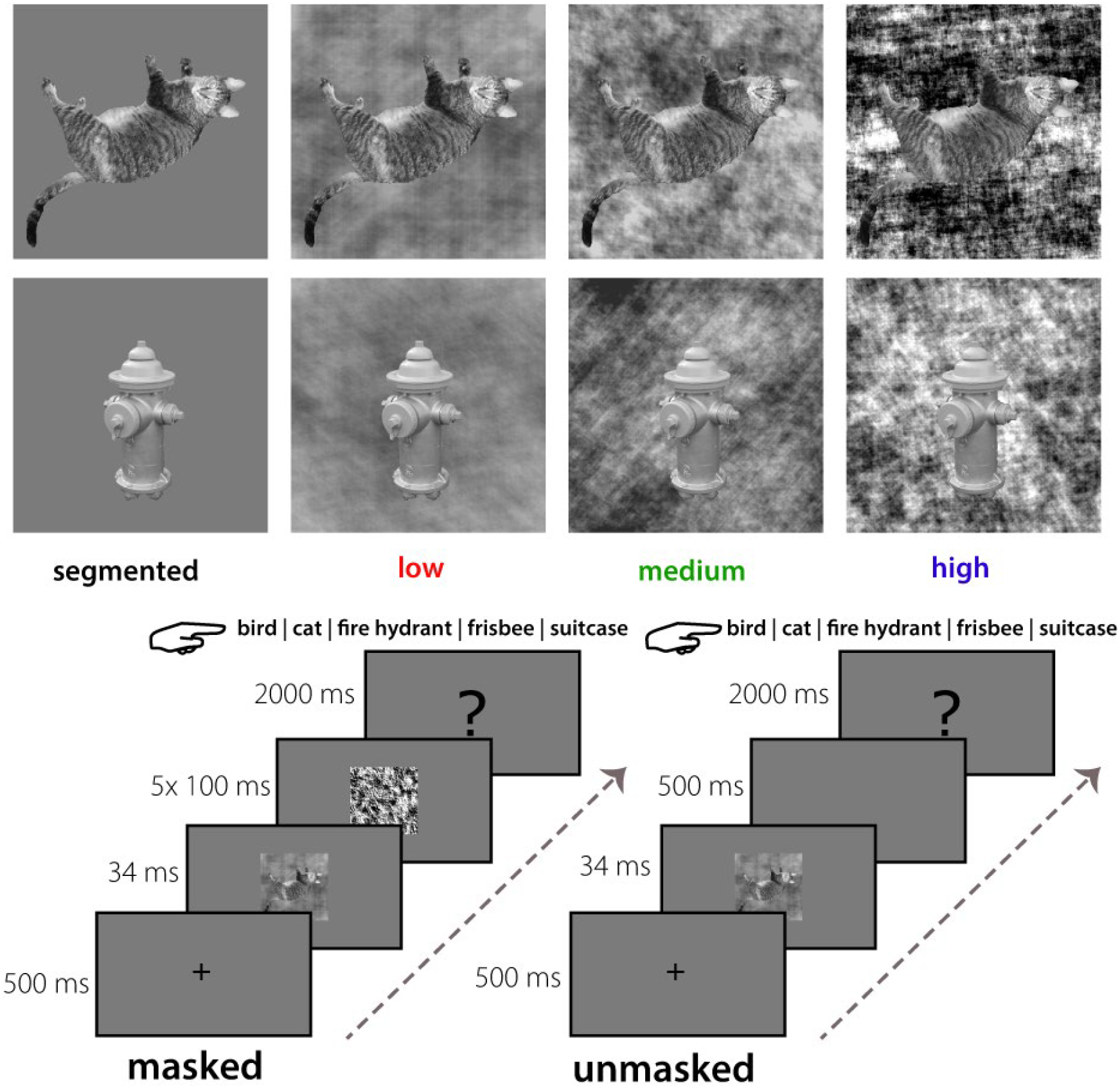
Stimuli and experimental paradigm. A) Exemplars of two categories (cat, fire hydrant) from each stimulus complexity condition. Backgrounds were either uniform (segmented; gray), or had low (red), medium (green) or high (blue) CE and SC values. B) Experimental design. On masked trials, the stimulus was followed by a dynamic mask (5×100 ms); on unmasked trials this was replaced by a blank screen (500 ms). Participants were asked to categorize the target object by pressing the corresponding button on the keyboard.

In half the trials, we impaired feedback activity with visual-masking. In addition to behavioral measures, we measured EEG responses to examine the time-course of visually evoked activity. Besides human participants, we also investigated recognition performance in Deep Convolutional Neural Networks (DCNNs), which received identical visual stimuli as our human participants, and performed a five-choice recognition task.

A convergence of results indicated that recurrent computations were critical for recognition of objects in complex environments, i.e. objects that were more difficult to segment from their background. First of all, behavioral results indicated poorer recognition performance for objects with more complex backgrounds, but only when feedback activity was disrupted by masking. Second, EEG measurements showed clear differences between complexity conditions in the ERPs around 200ms - a time point beyond the first feed-forward visual sweep of activity. Additionally, object category decoding based on the multivariate EEG patterns showed later decoding onsets for objects embedded in more complex backgrounds. This indicated that object representations for more complex backgrounds emerge later, compared to objects in more simple backgrounds. Finally, DCNN performance confirmed this interpretation; feed-forward networks showed a higher degree of impairment in recognition for objects in complex backgrounds compared to recurrent networks. Together, these results support the notion that recurrent computations drive figure-ground segmentation of objects in complex scenes.

## Materials and methods

### Subjects main experiment

Forty-two participants (32 females, 18-35 years old) took part in a first EEG experiment. Data from two participants were excluded from further analysis due to technical problems. We used this first dataset to perform exploratory analyses and optimize our analysis pipeline (Figure 2). Based on this dataset, we defined the time-windows (for further ERP analyses), electrode selection and preprocessing steps. To confirm our results on an independent dataset, another twenty participants (13 females, 18-35 years old) were measured. Data from one participant were excluded from ERP analyses, due to wrong placement of electrodes I1 and I2.

**Figure 2.**
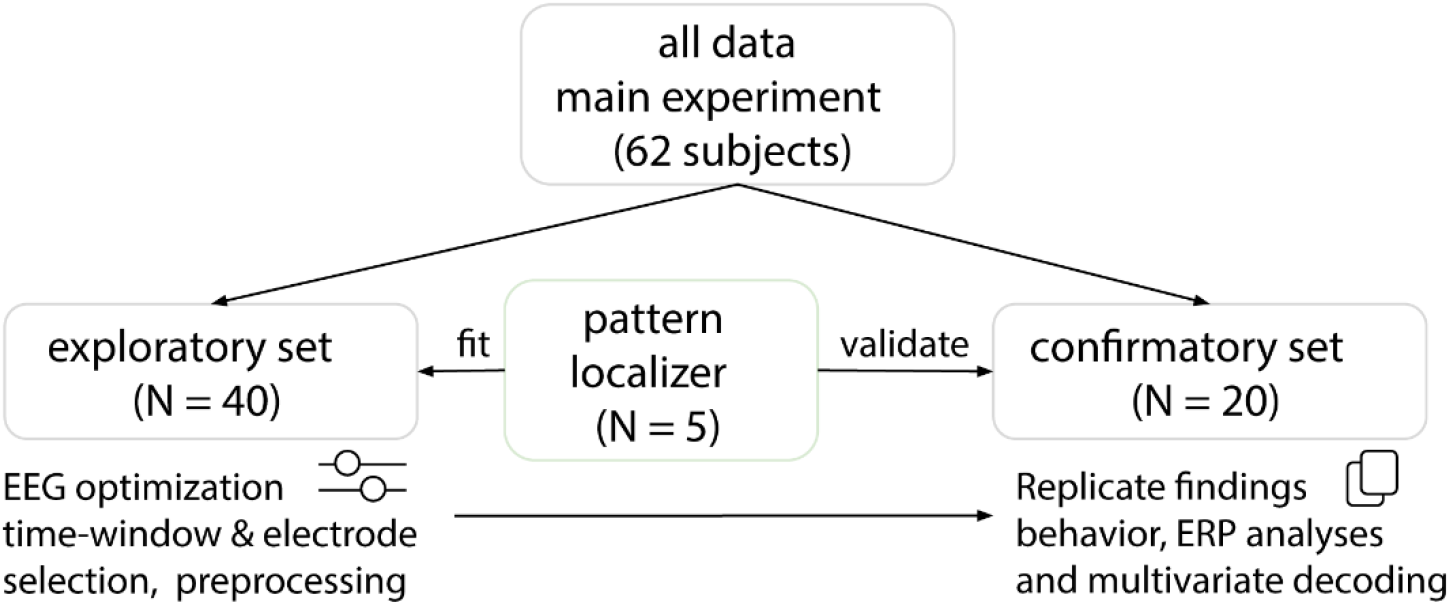
Experimental procedure. Sixty-two participants took part in the EEG experiment. Data from forty participants were used to perform exploratory analyses. The resulting data (twenty participants) were used to confirm our results. For the decoding analyses, five new participants took part in a separate experiment to characterize multivariate EEG activity patterns for the different object categories.

### Stimuli

Images of real-world scenes containing birds, cats, fire hydrants, frisbees or suitcases were selected from several online databases, including MS COCO (Lin et al., 2014), the SUN database (Xiao et al., 2010), Caltech-256 (Griffin et al., 2007), Open Images V4 (Kuznetsova et al., 2018) and LabelMe (Russell et al., 2008). These five categories were selected because a large selection of images was available in which the target object was clearly visible and not occluded. For each image, one CE and one SC value was computed using a simple visual model that simulates neuronal responses in one of the earliest stages of visual processing. Specifically, they are derived by averaging the simulated population response of LGN-like contrast filters across the visual scene (for a full description see Ghebreab et al. (2009), Scholte et al. (2009), Groen et al. (2013)). Computing these statistics for a large set of scenes results in a two-dimensional space in which sparse scenes with just a few scene elements separate from complex scenes with a lot of clutter and a high degree of fragmentation.

Together, CE and SC appear to provide information about the ‘segmentability’ of a scene (Groen et al., 2013, 2018). High CE/SC values correspond with images that contain many edges that are distributed in an uncorrelated manner, resulting in an inherently low figure-ground segmentation. Relatively low CE/SC values on the other hand correspond with a homogenous image containing few edges, resulting in an inherently high figure-ground segmentation (Figure 1). Each object was segmented from their real-world scene background and superimposed on three categories of phase scrambled versions of the real-world scenes.

This corresponded with low, medium and high complexity scenes. Additionally, the segmented object was also presented on a uniform gray background as the segmented condition (Figure 1). For each object category eight low CE/SC, eight medium CE/SC and eight high CE/SC images were selected, using the cut-off values from Groen et al. (2018), resulting in 24 images for each object category and 120 images in total. Importantly, each object was presented in all conditions, allowing us to attribute the effect to the complexity (i.e. segmentability) of each trial, and rule out any object-specific effects.

### Experimental design

Participants performed a 5-choice categorization task (Figure 1), differentiating images containing cats, birds, fire hydrants, frisbees and suitcases as accurately as possible. Participants indicated their response using five keyboard buttons corresponding to the different categories. Images were presented in a randomized sequence, for a duration of 34 ms. Stimuli were presented at eye-level, in the center of a 23-inch ASUS TFT-LCD display, with a spatial resolution of 1920*1080 pixels, at a refresh rate of 60 Hz. Participants were seated approximately 70 cm from the screen, such that stimuli subtended a 6.9° visual angle. The object recognition task was programmed in- and performed using Presentation (Version 18.0, Neurobehavioral Systems, Inc., Berkeley, CA, www.neurobs.com). The experiment consisted of 960 trials in total, of which 480 were backward masked trials and 480 were unmasked trials, randomly divided into eight blocks of 120 trials for each participant. After each block, participants took a short break. The beginning of each trial consisted of a 500 ms fixation period where participants focused their gaze on a fixation cross at the centre of the screen. In the unmasked trials, stimuli were followed by a blank screen for 500 ms and then a response screen for 2000 ms. In order to disrupt recurrent processes (Breitmeyer and Ogmen, 2000; Lamme et al., 2002; Fahrenfort et al., 2007), in the masked trials, five randomly chosen phase-scrambled masks were presented sequentially for 500 ms. The first mask was presented immediately after stimulus presentation, each mask was presented for 100 ms (Figure 1). The ambient illumination in the room was kept constant across different participants.

### Subjects pattern localizer

Five new participants took part in a separate experiment to characterize multivariate EEG activity patterns for the different object categories. For this experiment, we measured EEG activity while participants viewed the original experimental stimuli followed by a word (noun). Participants were asked to only press the button when the image and the noun did not match to ensure attention (responses were not analyzed). A classifier was trained on the EEG data from this experiment, and subsequently tested on the data from the main experiment using a cross-decoding approach. All participants had normal or corrected-to-normal vision, provided written informed consent and received monetary compensation or research credits for their participation. The ethics committee of the University of Amsterdam approved the experiment.

### Deep Convolutional Neural Networks (DCNNS)

First, to investigate the effect of recurrent connections, we tested different architectures from the CORnet model family (Kubilius et al., 2018); CORnet-Z (feed-forward), CORnet-R (recurrent) and CORnet-S (recurrent with skip connections). Then, to further evaluate the influence of network depth on scene segmentation, tests were conducted on three deep residual networks (ResNets; (He et al., 2016) with increasing number of layers; ResNet-10, ResNet-18 and Resnet-34. “Ultra-deep” residual networks are mathematically equivalent to a recurrent neural network unfolding over time, when the weights between their hidden layers are clamped (Liao and Poggio, 2016). This has led to the hypothesis that the additional layers function in a way that is similar to recurrent processing in the human visual system (Kar et al., 2019). Pre-trained networks were finetuned on images from the MSCoco database (Lin et al., 2014), using PyTorch (Paszke et al., 2017). After initialization of the pretrained network, the model’s weights were finetuned for our task, generating 5 probability outputs (for our 5 object categories). To obtain statistical results, we finetuned each network architecture ten different times.

### EEG data acquisition and preprocessing

EEG was recorded using a 64-channel Active Two EEG system (Biosemi Instrumentation, Amsterdam, The Netherlands, www.biosemi.com) at a 1024 Hz sample rate. As in previous studies investigating early visual processing (Groen et al., 2013, 2018), we used caps with an extended 10–10 layout modified with 2 additional occipital electrodes (I1 and I2, which replaced F5 and F6). Eye movements were recorded with additional electro-oculograms (vEOG and hEOG). Preprocessing was done using MNE software in Python (Gramfort et al., 2014) and included the following steps for the ERP analyses: 1) After importing, data were re-referenced to the average of two external electrodes placed on the mastoids. 2) A high-pass (0.1Hz, 0.1Hz transition band) and low-pass (30Hz, 7.5 Hz transition band) basic FIR filters were sequentially applied. 3) an Independent Component Analysis (ICA;(Vigario et al., 2000)) was run in order to identify and remove eye-blink and eye-movement related noise components (mean = 1.73 of first 25 components removed per participant). 4) epochs were extracted from −200 ms to 500 ms from stimulus onset. 5) trials were normalized by their 200 ms pre-stimulus baseline. 6) 5% of trials with the most extreme values within each condition were removed, keeping the number of trials within each condition equal. 7) data were transformed to current source density responses (Perrin et al., 1989).

### Statistical analysis: behavioral data

For human subjects, choice accuracy was computed for each condition in the masked and unmasked trials (Figure 3). Differences between the conditions were tested using two-factor (Scene complexity: segmented, low, med, high; Masking: masked, unmasked) repeated-measures ANOVAs. Significant main effects were followed up by post-hoc pairwise comparisons between conditions using Sidák multiple comparisons correction at α = 0.05. For DCNNs, a non-parametric Friedman test was used to differentiate accuracy across the different conditions (segmented, low, medium, high), followed by pairwise comparisons using a Mann-Whitney U test. Behavioral data were analyzed in Python using the following packages: Statsmodels, SciPy, NumPy, Pandas, (Jones et al., 2001; Oliphant, 2006; McKinney and Others, 2010; Seabold and Perktold, 2010).

**Figure 3.**
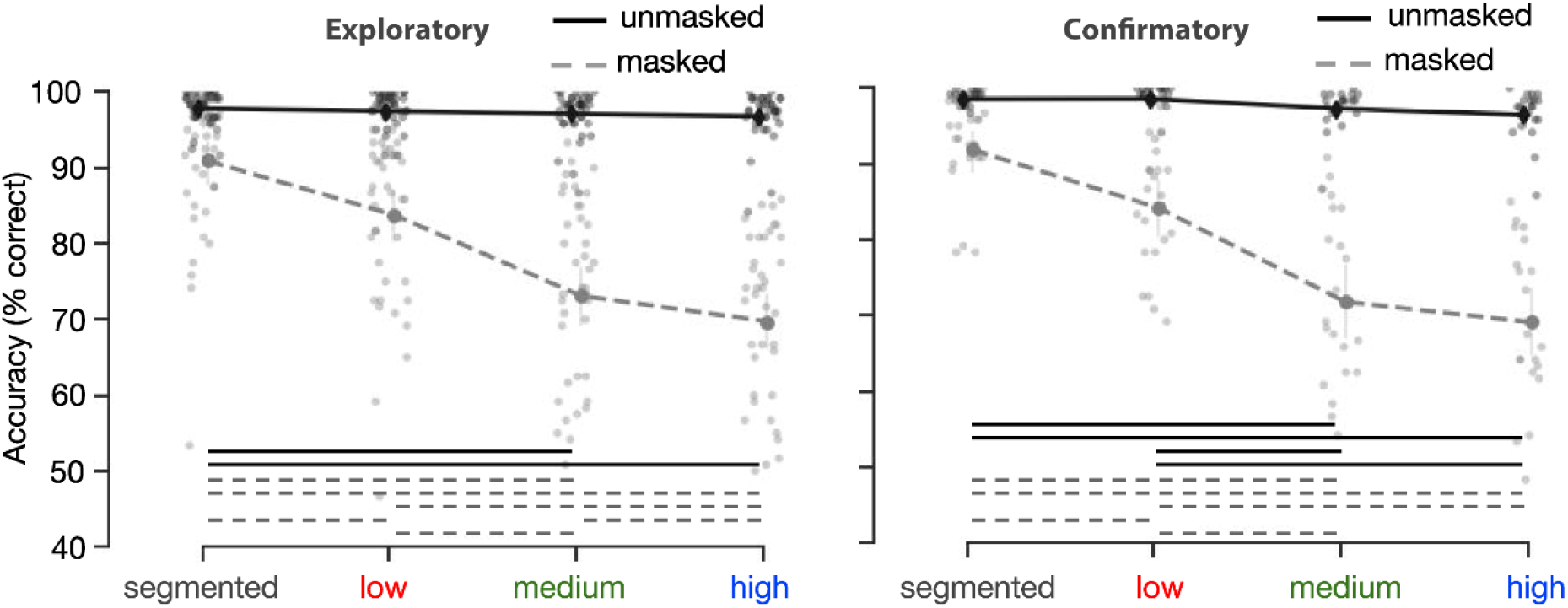
Human performance on the object recognition task. Performance (percentage correct) on the 5-option object recognition task. For masked trials, performance decreased with an increase in background complexity. The left panel shows results from the exploratory set (n=40), on the right results from the confirmatory set (N = 20) are plotted. Error bars represent the bootstrap 95% confidence interval, dots indicate the average performance of individual participants. Significant differences are indicated with a solid (unmasked) or dashed (masked) line.

### Statistical analysis: EEG - event related potentials

EEG analyses were carried out in Python, using the MNE software. For each participant, the difference in event-related potential (ERP) to scene complexity was computed within masked and unmasked conditions, pooled across occipital and peri-occipital electrodes (Oz, POz, O1, O2, PO3, PO4, PO7, PO8). This was done by subtracting the signal of each complexity condition (i.e. low, medium or high) from the segmented condition. Doing so enabled us to investigate differences between low, medium and high complex scenes regardless of masking effects. Based on the exploratory dataset, we established five time windows by performing t-tests on every time point for each condition and selecting windows in which the amplitude differed from zero for all complexity conditions (low, med, high). Then, a repeated measures ANOVA with factor background complexity (low, medium, high) and masking (masked, unmasked) was performed on the average activity in these established time windows.

### Statistical analysis: EEG - multivariate classification

The same preprocessing pipeline was used as for the ERP analyses. To evaluate how object category information in our EEG signal evolves over time, cross-decoding analyses were performed by training a Support Vector Machine (SVM) classifier on all trials from the pattern localizer experiment (performed by five different subjects) and testing it on each of the main experiment conditions. Object category classification was performed on a vector of EEG amplitudes across 22 electrodes, including occipital (I1, Iz, I2, O1, Oz, O2), peri-occipital (PO3, PO7, POz, PO4, PO8), and parietal (Pz, P1-P10) electrodes. EEG activity was standardized and averaged across the five time windows derived from the ERP analyses. Statistical significance was determined using a Wilcoxon signed-rank test, and corrected for multiple comparisons using a false discovery rate (FDR) of 0.05.

## Results

### Behavior

During the task, participants viewed images of objects placed on top of a gray (segmented), low, medium or high complexity background. On each trial, they indicated which object category the scene contained, using the corresponding keyboard buttons. In half of the trials, the target image was followed by a dynamic backward mask (5×100 ms); the other half of the trials was unmasked (Figure 1). Accuracy (percentage correct trials) was computed for each participant. A repeated measures ANOVA on the exploratory dataset (N = 40), with factors background (segmented, low, medium, high) and masking (masked, unmasked) indicated, apart from main effects, an interaction effect. Results indicated that masking impaired performance for objects presented on more complex backgrounds stronger than for less complex backgrounds (F(3,117) = 185.6748, *p* < .001). Post-hoc comparisons showed that for masked trials, accuracy decreased for both medium (t(39) = 2.88, *p*(Sidák-corrected) = 0.038) and high (t(39) = 3.84, *p*(Sidák-corrected) = 0.003) complexity condition compared to the low condition (all other *p* > .203). For unmasked trials, all conditions differed from each other, with an incremental decrease in accuracy for objects presented on more complex backgrounds.

Analysis of the confirmatory dataset (N = 20) indicated similarly, apart from the main effects, an interaction between masking and background complexity. For masked trials, there was a larger decrease in performance with an increase in background complexity, (F(3, 57) = 101.3338, *p* < .001). Post-hoc comparisons showed that for masked trials, accuracy decreased for both medium and high complexity conditions compared to the segmented (t(19) = 3.47, *p*(Sidák-corrected) = 0.003, (t(19) = 3.47, *p*(Sidák-corrected) = 0.003) and low conditions (t(19) = 4.23, *p*(Sidák-corrected) < .001, (t(19) = 4.31, *p*(Sidák-corrected) < .001). For unmasked trials, all conditions differed from each other with the exception of medium - high, with an incremental decrease in accuracy for objects presented on more complex backgrounds.

### Network performance

Next, we presented the same images to Deep Convolutional Neural Networks with different architectures. For the CORnets (Figure 4, left panel), a non-parametric Friedman test differentiated accuracy across the different conditions (segmented, low, medium, high) for all architectures, Friedman’s Q(3) = 27.8400; 24.7576; 26.4687 for CORnet-Z, -RT -S respectively, all *p* < .001. A Mann-Whitney U test on the difference in performance between segmented and high complexity trials indicated a smaller decrease in performance for CORnet-S compared to CORnet-Z (Mann–Whitney U = 100.0, n1 = n2 = 10, *p* < .001, two-tailed). For the ResNets (Figure 4, right panel), a non-parametric Friedman test differentiated accuracy across the different conditions for ResNet-10 and ResNet-18, Friedman’s Q (3) = 23.9053; 22.9468, for ResNet-10 and ResNet-18 respectively, both *p* < .001. A Mann-Whitney U test on the difference in performance between segmented and high complexity trials indicated a smaller decrease in performance for ResNet-34 compared to ResNet-10 (Mann– Whitney U = 99.0, n1 = n2 = 10, *p* < .001, two-tailed). Overall, in line with human performance, results indicated a higher degree of impairment in recognition for objects in complex backgrounds for feed-forward or more shallow networks, compared to recurrent or deeper networks.

**Figure 4.**
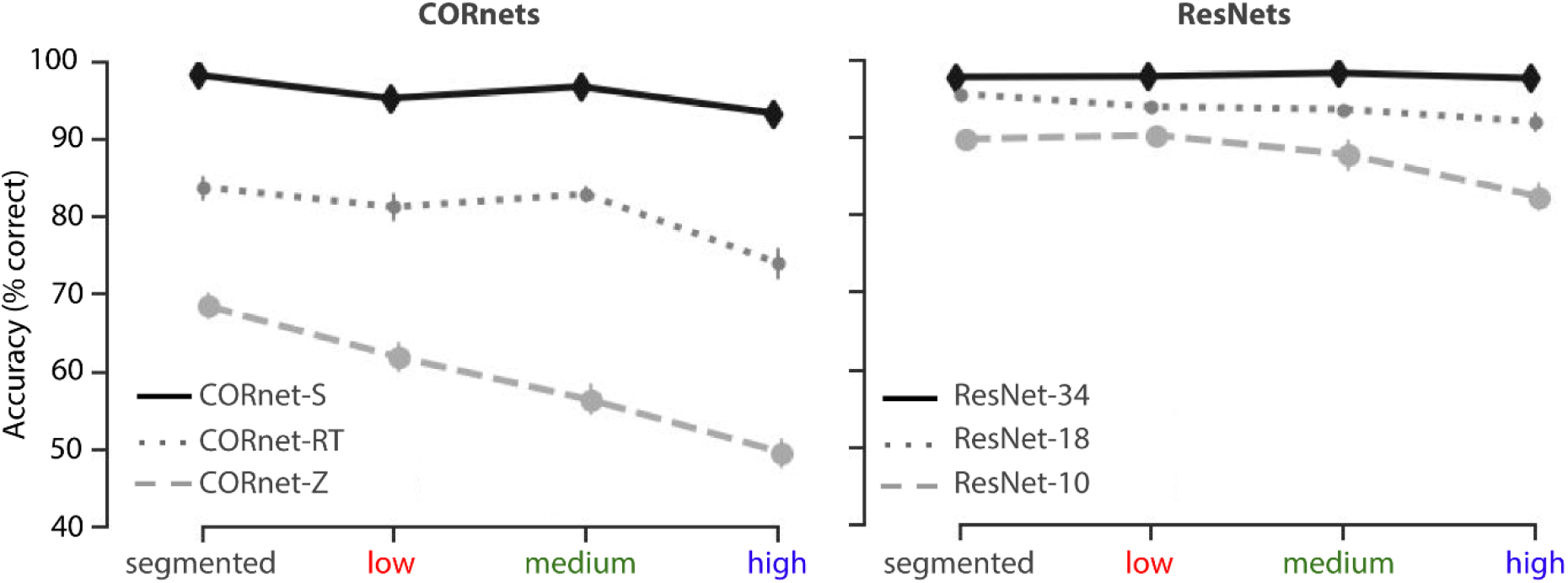
Deep Convolutional Neural Network performance on the object recognition task. Performance (percentage correct) on the 5-option object recognition task. Networks were finetuned on the 5 target categories, top-1 accuracy was computed. For the CORnets (left panel) performance of the feedforward architecture decreased with an increase in background complexity. For recurrent architectures, this decrease was less prominent. For CORnet-S, there was no difference between conditions. Error bars represent the bootstrap 95% confidence interval.

### EEG - event related potentials

To investigate the time-course of figure-ground segmentation in visual cortex, evoked responses to the masked and unmasked scenes were pooled across occipital and peri-occipital electrodes (Oz, POz, O1, O2, PO3, PO4, PO7, PO8), for each condition. Difference waves were generated by subtracting the signal of each condition from the segmented condition (Figure 5B/E). Doing so enabled us to eliminate the effect of masking on the EEG signal, and to investigate differences between low, medium and high complex scenes. For each participant, data was averaged across five time windows based on analyses on the exploratory dataset (see Materials and methods).

**Figure 5.**
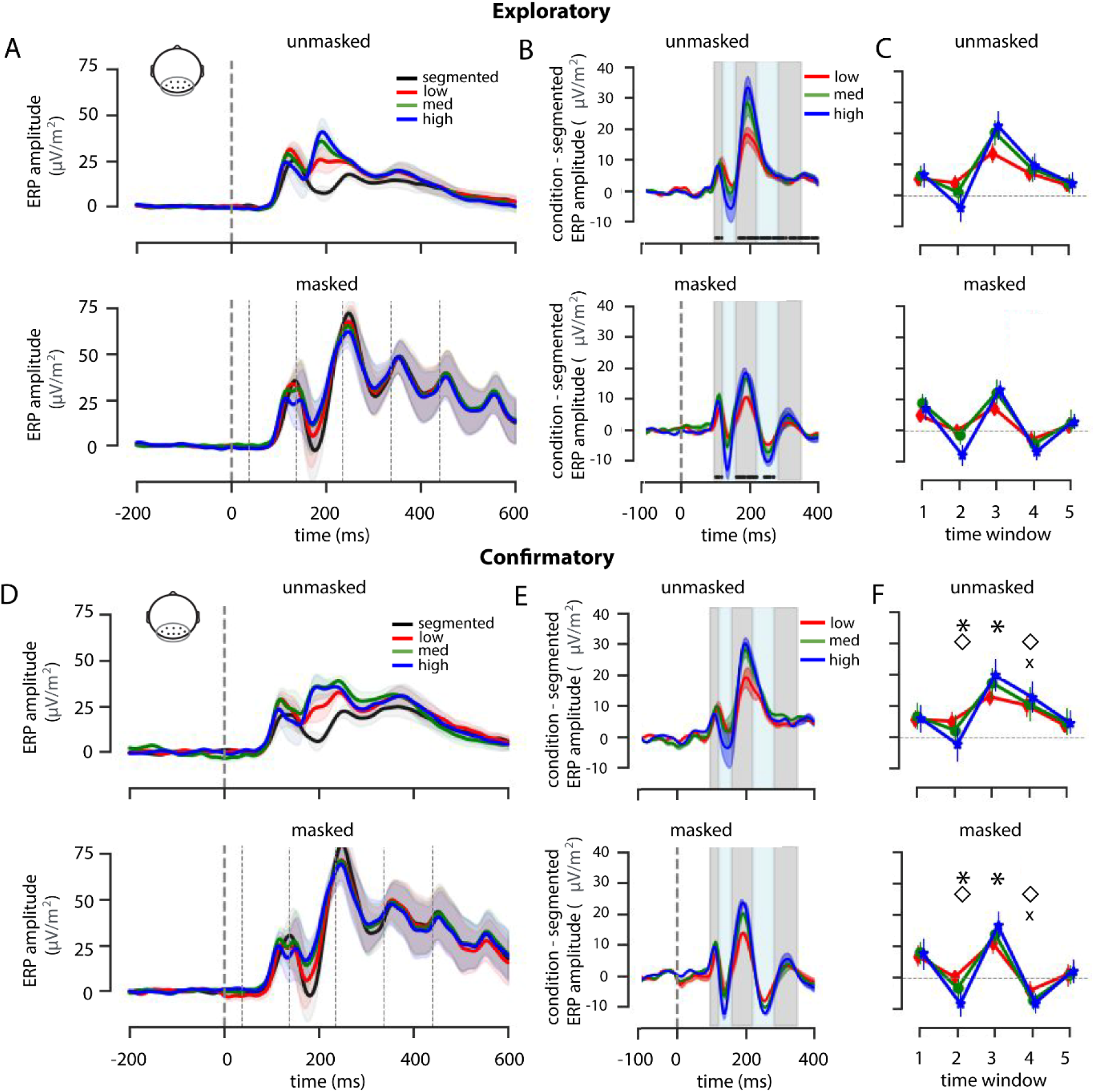
ERP results. **A)** Average ERP amplitude for segmented, low, medium and high complexity scenes for an occipital-peri-occipital pooling of EEG channels (Oz, POz, O1, O2, PO3, PO4, PO7, PO8) for masked and unmasked trials. Shaded regions indicate SEM across participants. Mask onsets are indicated with thin dashed lines (bottom panel only) **B)** Difference waves were generated by subtracting the signal of each condition from the segmented condition. Five time windows were determined by performing t-tests on every time point for each condition and selecting windows in which the amplitude differed from zero for all complexity conditions. Significant timepoints are indicated with a black dot above the x-axis. **C)** Based on significant timepoints in the exploratory dataset, five time-windows were defined: 92-115 ms; 120-150 ms; 155-217 ms; 221-275 ms; 279-345 ms). **D/E/F)** Analyses repeated for the confirmatory dataset. Symbol markers indicate main or interaction effects, asterisk: main effect of Complexity; diamond: main effect of Masking; plus: interaction effect.

For every time window, a Repeated Measures ANOVA was performed on the average EEG amplitude of the difference waves, with Complexity (low, med, high) and Masking (masked, unmasked) as within subject factors. As preprocessing procedure and time point selection were based on t-tests on the exploratory set, we do not report subsequent Repeated Measures ANOVA for this dataset. Results on the confirmatory dataset (Figure 5D/E/F) showed no main-or interaction effects in the first time-window (92-115 ms; Figure 5F). Critically, differences between Complexity conditions only emerged in time-window 2 and 3 (120-150 ms: F(36) = 22.87, *η^2par^* = .56, *p* < .001; 155-217 ms: F(36) = 24.21, *η^2par^* = .57, *p* < .001), suggesting a differential contribution of recurrent processing to object recognition in varying complexity scenes. In time-window 2, there was a main effect of Masking (F(18) = 5.38, *η^2par^* = .576, *p* = .03. Only in time window 4 (221-275 ms), an interaction effect of Masking and Complexity, F(18) = 59.60, *η^2par^* = .07, *p* < .001 started to emerge.

### EEG multivariate classification

To further investigate the representational dynamics of object recognition under different complexity conditions, multivariate decoding analyses were performed on the averaged activity in the five time windows (Figure 6). To control for response-related activity (keyboard buttons were fixed across the task), a cross-decoding analysis was performed, by training the classifier on all trials from an independent pattern localizer experiment, and testing it on each of the main experiment conditions (see Methods for details). For unmasked trials, a Wilcoxon signed-rank test on the exploratory dataset indicated successful decoding for segmented trials in all five time windows (Z = 100, *p* < 0.001; Z = 89, *p* < 0.001; Z = 30, *p* < 0.001; Z = 131, *p* < 0.001; Z = 141, *p* < 0.001) and low trials in the first three time windows (92-115 ms; 120-150 ms; 155-217 ms; Z = 198, *p* = 0.007; Z = 82, *p* < 0.001; Z = 61, *p* < 0.001). For objects on medium complex background, successful above-chance decoding emerged slightly later, in time-windows 2 and 3 (Z = 200, *p* = 0.012; Z = 110, *p* < 0.001). For objects on high complex background, there was successful decoding in time window 3, Z=216, p = 0.045. For masked trials, there was successful decoding for the segmented objects in time-windows 1, 3 and 4, Z = 113, *p* < 0.001; Z = 183, *p* = 0.004; Z = 186, *p* = 0.004, followed by later additional decoding of low (155-217 ms), Z = 138, *p* = 0.001, and high (221-275 ms) complexity trials, Z = 157, *p* = 0.003. There were no significant time windows for medium complexity trials. All p-values reported were corrected by FDR = 0.05.

**Figure 6.**
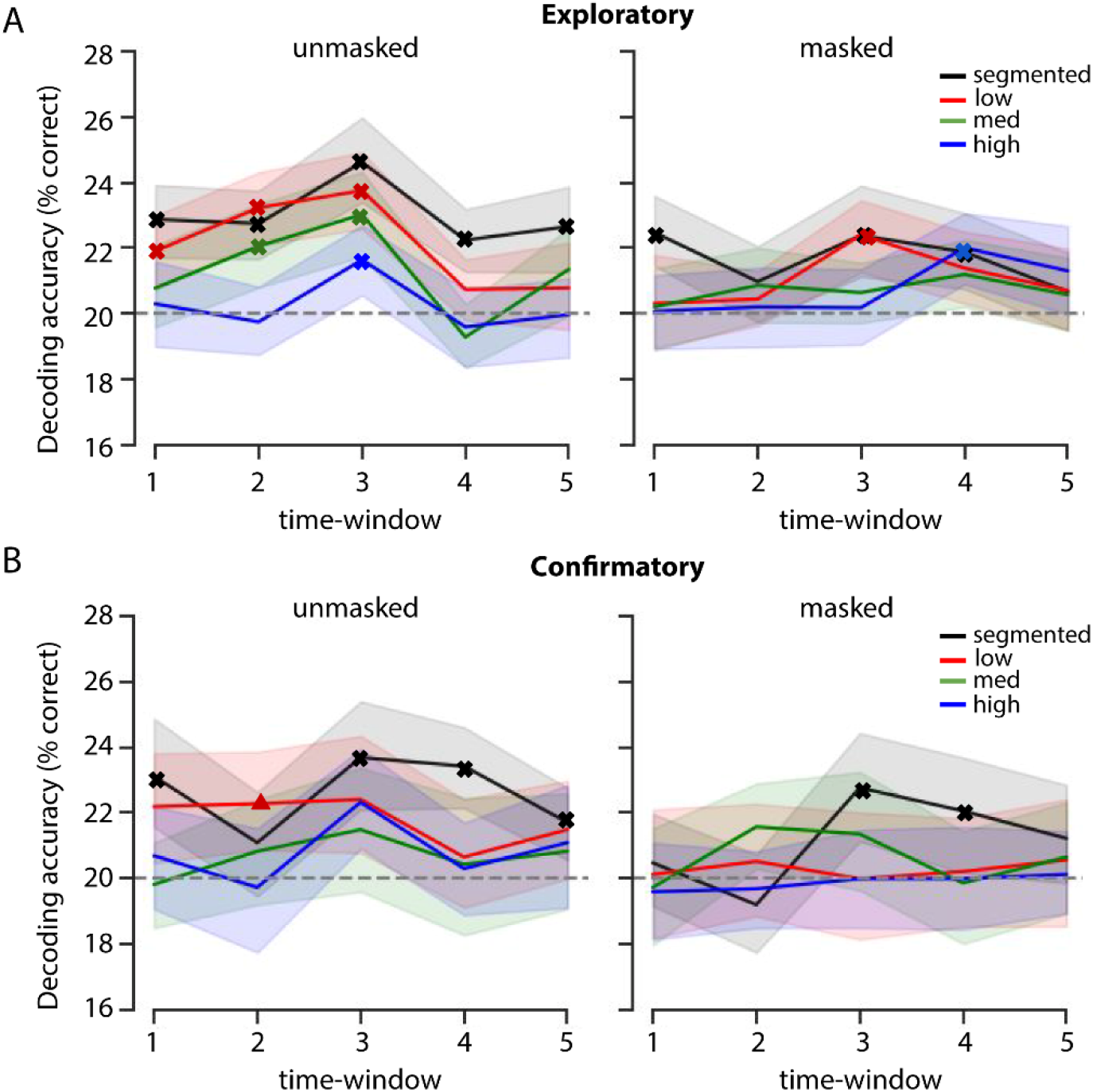
Cross-decoding results using the pattern localizer. Decoding object category in the EEG signal for the A) exploratory dataset, and the B) Confirmatory dataset, for masked and unmasked trials with varying complexity in the five time windows. The dotted line represents the 20% chance-level, shaded error bars represent the bootstrap 95% confidence interval. Results from the Wilcoxon signed-rank test are indicated with a bold **x** (corrected for multiple comparisons using a false discovery rate of 0.05).

Finally, we aimed to replicate these findings in the confirmatory dataset (N = 20). Overall, results indicated fewer instances of successful object decoding, and if present, slightly delayed compared to the exploratory set. For unmasked trials, results from the Wilcoxon Signed-Ranks test indicated successful decoding for segmented trials (92-115 ms; 155-217 ms; 221-275 ms; 279-245 ms), Z = 27, *p* = 0.006; Z = 18, *p* = 0.003; Z = 0, *p* < 0.001; Z = 35, *p* = 0.011. There were no other significant time windows from other unmasked conditions. For masked conditions, there was also only significant decoding in segmented trials, specifically in time window 3 and 4 (155-217 ms; 221-275 ms), Z = 36, *p* = 0.031; Z = 38, *p* = 0.031.

Overall, these findings showed that different objects evoked reliably different sensor patterns when presented in isolation or in ‘simple’ environments, within the first feed-forward sweep of visual information processing. Additionally, results indicated decreased and later decoding for objects embedded in more complex backgrounds, suggesting that object representations for objects on complex backgrounds emerge later. Finally, these findings also indicate that the object category representations generalized across tasks and participants.

## Discussion

This study systematically investigated whether recurrent processing is required for figure-ground segmentation during object recognition. A converging set of behavioral, EEG and computational modelling results indicate that recurrent computations are required for figure-ground segmentation of objects in complex scenes. These findings are consistent with previous findings showing enhanced feedback for complex scenes (Groen et al., 2018), and visual backward masking being more effective for images that were ‘more difficult to segment’ (Koivisto et al., 2014). We interpret these results as showing that figure-ground segmentation, driven by recurrent processing, is not necessary for object recognition in simple scenes but it is for more complex scenes.

### Effects of scene complexity using artificial backgrounds

In an earlier study, using natural scenes, we already showed that feedback was selectively enhanced for high complexity scenes, during an animal detection task. While there are numerous reasons for using naturalistic scenes (Felsen and Dan, 2005; Felsen et al., 2005; Talebi and Baker, 2012), it is difficult to do controlled experiments with them because they vary in many (unknown) dimensions. Additionally, SC and CE (measures of scene complexity) could correlate with other contextual factors in the scene (e.g. SC correlates with perception of naturalness of a scene (Groen et al., 2013)), and could be used as diagnostic information for the detection of an animal. Additionally, previous research has shown that natural scenes and scene structure can facilitate object recognition (Davenport and Potter, 2004; Neider and Zelinsky, 2006; Kaiser and Cichy, 2018). Results from the current study, using artificial backgrounds of varying complexity, replicate earlier findings while allowing us to attribute the effects to SC and CE, and the subsequent effect on segmentability. A limitation of any experiment with artificially generated (or artificially embedded) images is that it’s not clear whether our findings will generalize to ‘real images’ that have not been manipulated in any way. Together with the previous findings, however, our results corroborate the idea that more extensive processing (possibly in the form of recurrent computations) is required for object recognition in more complex, natural environments (Groen et al., 2018; Rajaei et al., 2019).

### Time course of object recognition

Based on the data from the exploratory dataset (N = 40), we selected five time windows in the ERPs to test our hypotheses on the confirmatory dataset. For our occipital-peri-occipital pooling, we expected the first feed-forward sweep to be unaffected by scene complexity (i.e. low, med, high). Indeed, amplitudes of the difference waves (complexity condition - segmented ERP amplitudes) averaged across the selected time windows indicated no influence of masking or scene complexity early in time (92-115 ms). The observation that all three difference waves deviated from zero, however, indicates that there was an effect of segmentation. In this early time window, background presence thus seems to be more important than the complexity of the background. This difference could be attributed to the detection of additional low-level features in the low, medium and high complexity condition, activating a larger set of neurons that participate in the first feed-forward sweep (Lamme and Roelfsema, 2000). In the second and third time window (120-217 ms), differences between the complexity conditions emerged. We interpret these differences as reflecting the increasing need for recurrent processes when backgrounds are more complex.

Our results are generally consistent with prior work investigating the time course of visual processing of objects under more or less challenging conditions (DiCarlo and Cox, 2007; Cichy et al., 2014; Contini et al., 2017; Tang et al., 2018; Rajaei et al., 2019). In line with multiple earlier studies, masking left the early evoked neural activity (<120 ms) relatively intact, whereas the neural activity after ∼150 ms was decreased (Lamme and Roelfsema, 2000; Lamme et al., 2002; Del Cul et al., 2007; Fahrenfort et al., 2007; Boehler et al., 2008; Koivisto and Revonsuo, 2010).

Decoding results corroborated these findings, showing decreased or delayed decoding onsets for objects embedded in more complex backgrounds, suggesting that object representations for those images emerge later. Additionally, when recurrent processing was impaired using backward masking, only objects presented in isolation or in ‘simple’ environments evoked reliably different sensor patterns that our classifiers were able to pick up (Figure 5 and 6).

### Influence of masking on behavior

Based on the strong interaction effect on behavior, it is tempting to conclude that complexity significantly increases the effect of masking on recognition accuracy. However, performance on all unmasked trials was virtually perfect (96-97%) raising concerns about ceiling effects obscuring the actual variation between these conditions (Uttl, 2005). Therefore, although masked stimuli show a decrease in performance along increases in complexity; based on the current findings we cannot conclude that this is because of masking (i.e. reducing recurrent processes). While we do not claim that unmasked segmented, low, medium, or high images are equally difficult or processed in the same way (we actually argue for the opposite), our results show that apparently the brain is capable of arriving at the correct answer with enough time. It is hard to come up with an alternative (more difficult) task without affecting our experimental design and subsequent visual processing (e.g. stimulus degradation generally affects low-level complexity; reducing object size or varying object location creates a visual search task that could benefit from spatial layout properties). Combined fMRI and EEG results from an earlier study already showed that for complex scenes only, early visual areas were selectively engaged by means of a feedback signal (Groen et al., 2018). Here, using controlled stimuli and backward masking, we replicate and expand on these findings. Importantly, results from both EEG and deep convolutional neural networks support the notion that recurrent computations drive figure-ground segmentation of objects in complex scenes.

### Consistency of object decoding results

In the exploratory set, results from the multivariate decoding analyses indicated early above chance decoding for ‘simple’ scenes (segmented and low) in both unmasked and masked trials. For more complex scenes decoding emerged later (medium) or was absent (high) for unmasked trials. In the confirmatory set, however, there were fewer instances of successful object decoding, and if present, successful decoding was delayed. A likely explanation for this finding could be that the sample size in the confirmatory dataset was inadequate for the chosen multivariate decoding analyses, resulting in reduced statistical power. A simulation analysis on the exploratory set, in which we randomly selected 20 participants (repeated 1000 times) indicated reduced decoding accuracy, similar to our confirmatory results. Our choice for the number of participants in the confirmatory dataset thus does not seem to be sufficient (Supplementary Figure S1).

### Probing cognition with Deep Convolutional Neural Networks

One way to understand how the human visual system processes visual information involves building computational models that account for human-level performance under different conditions. Here we used Deep Convolutional Neural Networks, because they show remarkable performance on both object and scene recognition (e.g. (Russakovsky et al., 2015; He et al., 2016). While we certainly do not aim to claim that DCNNs are identical to the human brain, we argue that studying how performance of different architectures compares to human behaviour *could* be informative about the type of computations that are underlying this behavior (Cichy and Kaiser, 2019). In the current study, it provides an additional test for the involvement of recurrent connections. Comparing the (behavioral) results of DCNNs with findings in humans, our study adds to a growing realization that more extensive processing, in the form of recurrent computations, is required for object recognition in more complex, natural environments (Groen et al., 2018; Tang et al., 2018; Kar et al., 2019; Rajaei et al., 2019).

## Conclusion

Results from the current study show that how object recognition is resolved depends on the context in which the target object appears: for objects presented in isolation or in ‘simple’ environments, object recognition appears to be dependent on the object itself, resulting in a problem that can likely be solved within the first feed-forward sweep of visual information processing on the basis of unbound features (Crouzet and Serre, 2011). When the environment is more complex, recurrent processing is necessary to group the elements that belong to the object and segregate them from the background.

## Supplementary Figures

**Supplementary Figure S1.**
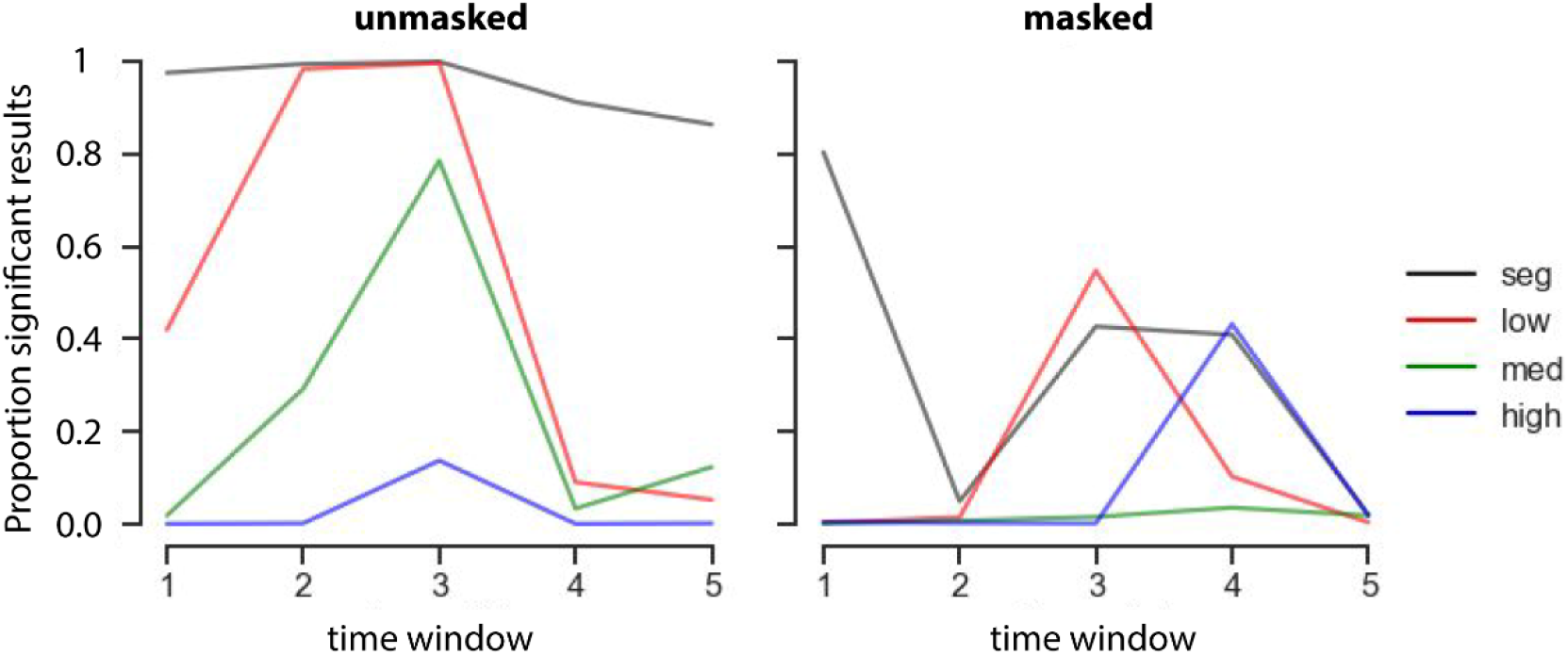
Simulation analysis on the exploratory set. Random selection of 20 participants (repeated 1000 times) indicated reduced chances of finding significant decoding results. Plotted are the proportion (number of instances divided by 1000) in which the results remained significant.

## Acknowledgements

This work is supported by and Advanced Investigator Grant from the European Research Council (ERC) to Edward de Haan (<339374) and an Interdisciplinary Doctorate Agreement from the University of Amsterdam to H. Steven Scholte and Jessica Loke.

## Data and code availability

Data and code to reproduce the analyses in this article are available at https://github.com/noorseijdel/2020_EEG_figureground

